# Learning Protein-Protein Interaction Interfaces from Multimodal Representations

**DOI:** 10.64898/2026.05.29.728739

**Authors:** Yuri Gardinazzi, Edith Natalia Villegas Garcia, Sergio Senci, Davide Di Vora, Antonio Feltrin, Francesca Cuturello

## Abstract

**Background:** Protein-protein interactions govern essential cellular processes, making the identification of interface residues critical for protein engineering and molecular design. Although complex deep learning algorithms like AlphaFold3 generate full atom resolution assemblies, they are not optimized for residue-level interface identification. Existing methods face inherent limitations: sequence-based models lack explicit 3D context, while structure-based approaches can overlook evolutionary signals.

**Results:** By combining evolutionary context with 3D geometry, the multimodal protein language model ESM3 offers the ideal foundation for residue-level interface classification. This results in the ESM3-PPISites model, which outperforms existing state-of-the-art methods across diverse training and benchmark sets. Generalization is assessed through stringent sequence- and structure-based redundancy filtering to ensure evaluation on previously unseen proteins. Notably, finetuning the open-weight ESM3 architecture meaningfully narrows the performance gap with its proprietary full-scale counterpart. Furthermore, integrating predicted interaction sites as spatial restraints into HADDOCK significantly improves the recovery of near-native binding poses while reducing computational search time by an order of magnitude compared with blind docking.

**Conclusions:** ESM3-PPISites enables accurate interfaces identification without relying on complex structural inputs at inference time, establishing a reliable and computationally efficient strategy for mapping protein-protein interaction sites.

## 1 Introduction

Protein-protein interactions (PPI) underlie most cellular processes, ranging from signal transduction and immune responses to macromolecular assembly. Decades of structural biology studies have established that PPI interfaces are shaped by physical and biochemical constraints such as accessible surface area burial, tight packing, specific residue propensities, and localized energy hot spots [1–9]. These molecular signatures reflect a functional surface adaptation depending on the interaction’s biological context, transitioning from large hydrophobic surface areas to smaller, polar interfaces in transient complexes [10]. Because the physicochemical patterns driving interface formation are complex and context-dependent, deciphering them requires advanced computational frameworks. In the era of full-atom structure predictors like AlphaFold3 [11], a crucial gap remains: structure predictors are neither designed nor optimized for partner-independent interface identification, whereas residue-level interface propensities can provide actionable guidance for site-directed mutagenesis and epitope mapping directly from individual protein sequences, without requiring a structural model of the interacting complex. Over the years, machine learning predictors have evolved along two primary input modalities: sequence-based and structure-based approaches [12–14]. Early sequence-based methods relied on traditional classifiers [15–26], but the domain was rapidly transformed by deep learning architectures [27–31]. Structure-based methods moved beyond hand-crafted features and biophysical potentials [32–35] toward deep architectures operating directly on 3D coordinates. In particular, Graph Neural Networks emerged as a natural choice to model protein structures [36–40], which subsequently paved the way for Geometric Deep Learning techniques to directly capture 3D spatial rotational and translational symmetries [41–45]. A significant advance emerged with Protein Language Models (PLMs) [46–51], a specialized class of self-supervised models that learn high-dimensional representations from vast sequence databases to capture complex evolutionary patterns. Through extensive pre-training, these foundational models are well-suited for finetuning on downstream tasks, such as identifying residues with strong interaction propensity [52–55]. (However, most PLMs are trained on sequence alone and capture structural information only implicitly. Recent multimodal models such as ESM3 jointly encode sequence, structure and function, natively bridging evolutionary signals with 3D structural context and thus offering a natural paradigm for partner-independent interface identification.) By natively bridging evolutionary sequence signals with 3D structural context, multimodal PLMs offer an ideal framework for partner-independent interface identification.

Here, we present ESM3-PPISites, a supervised residue-level classification model that leverages multimodal ESM3 representations [56] coupled with a lightweight dense classifier to identify protein interaction sites. Across diverse benchmarks, ESM3-PPISites significantly outperforms state-of-the-art methodologies. To prevent data leakage, sequence and structural homology biases were strictly eliminated during benchmarking. Despite marked differences in training data scale and composition, the model maintains high predictive fidelity across unseen protein families by effectively exploiting structural diversity. Beyond standalone prediction, integrating these predicted sites as spatial restraints into HADDOCK [57] substantially restricts the conformational search space, accelerating dockings while improving the recovery of near-native complex structures (Fig. 1). Altogether, ESM3-PPISites establishes a fast and reliable approach for interface mapping.

**Fig. 1.**
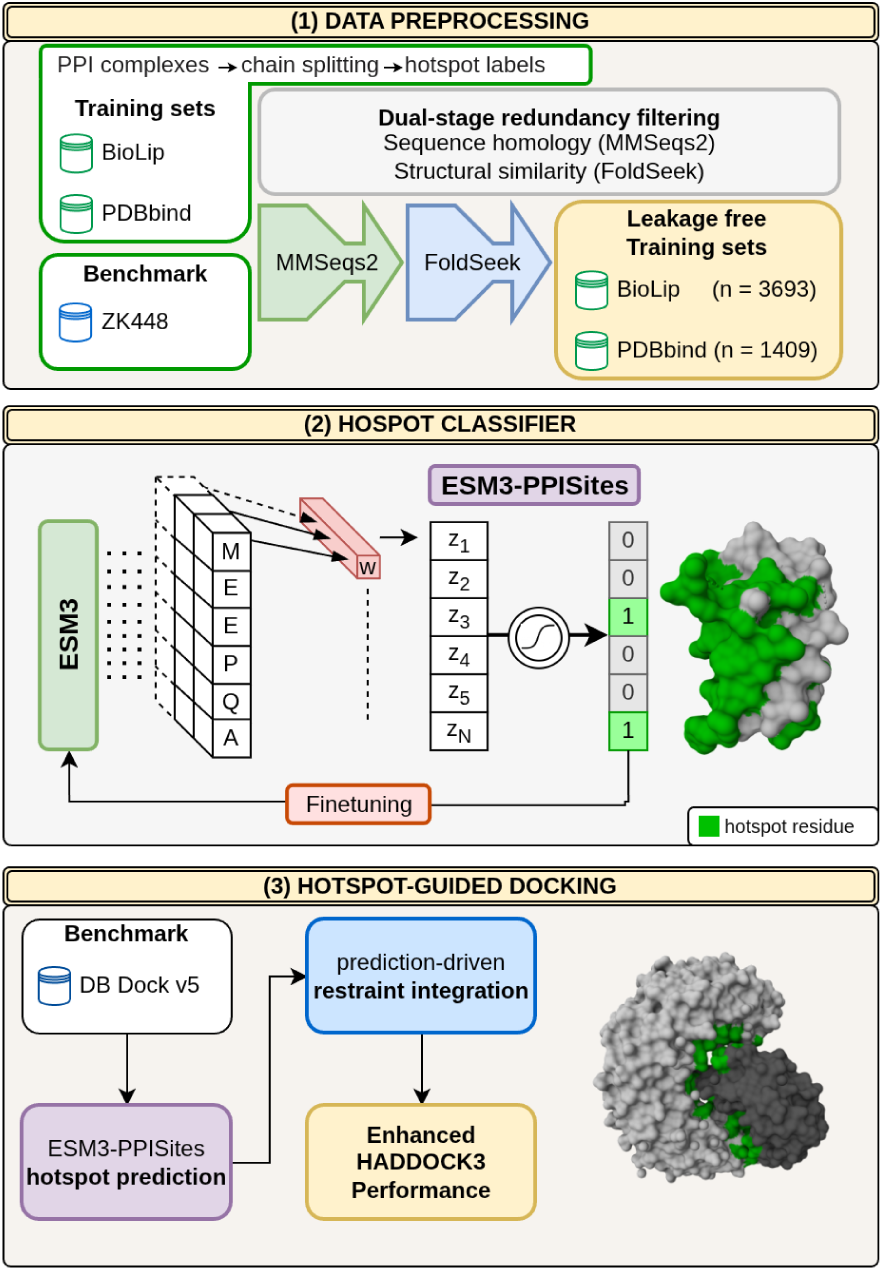
Schematic workflow of ESM3-PPISites. (1) Leakage-free dataset construction; (2) adaptation of ESM3 for PPI sites classification; (3) integration of predictions as restraints in HADDOCK3 docking.

## 2 Methods

### 2.1 Datasets and Curation

We use two data sources for model training: **BioLiP** [58], from which we adopt the provided validation set, and an in-house dataset derived from **PDBbind** v2020 [59, 60]. For the latter, we select an initial pool of 15,000 protein chains, excluding complexes involving nucleic acids or peptides, and cluster the sequences using MMseqs2 at 25% sequence similarity. To construct the validation set, we randomly sample 10% of the sequences from the smallest clusters. This strategy promotes sequence diversity and reduces redundancy, providing a heterogeneous subset for grid-search hyperparameter tuning. Interface labels are assigned at the residue level using the reference PDB complexes. Two residues from different chains are considered to be in contact if the distance between any pair of their atoms is below a threshold defined as 0.5 Å plus the sum of the corresponding van der Waals radii [12, 58]. Residues involved in at least one such contact are labelled as interface residues. To assess generalization on an independent benchmark, we use the **ZK448 dataset** [12], originally compiled by Zhang and Kurgan and retrieved from the PIPENN method repository [29]. Considered a gold-standard benchmark for PPI interface prediction, ZK448 comprises 336 non-redundant protein chains extracted from high-resolution hetero-complexes, with a maximum pairwise sequence identity of 25%. To guarantee a rigorous assessment of model generalization and prevent data leakage, we implement a dual-stage redundancy filtering protocol between the training candidate sets and the ZK448 benchmark (Figure 2):

1. **Sequence-Level Filtering:** MMseqs2 [61] is employed to exclude any training sequence sharing more than 25% identity with any entry in the ZK448 benchmark set.
2. **Structural-Level Filtering:** Foldseek [62] is subsequently applied to the structures to remove remaining homologs. Training proteins exhibiting an *E*-value below 10*^−^*^3^ and a target- or query-normalized TM-score exceeding 0.5 relative to any benchmark entry are strictly excluded.

**Fig. 2.**
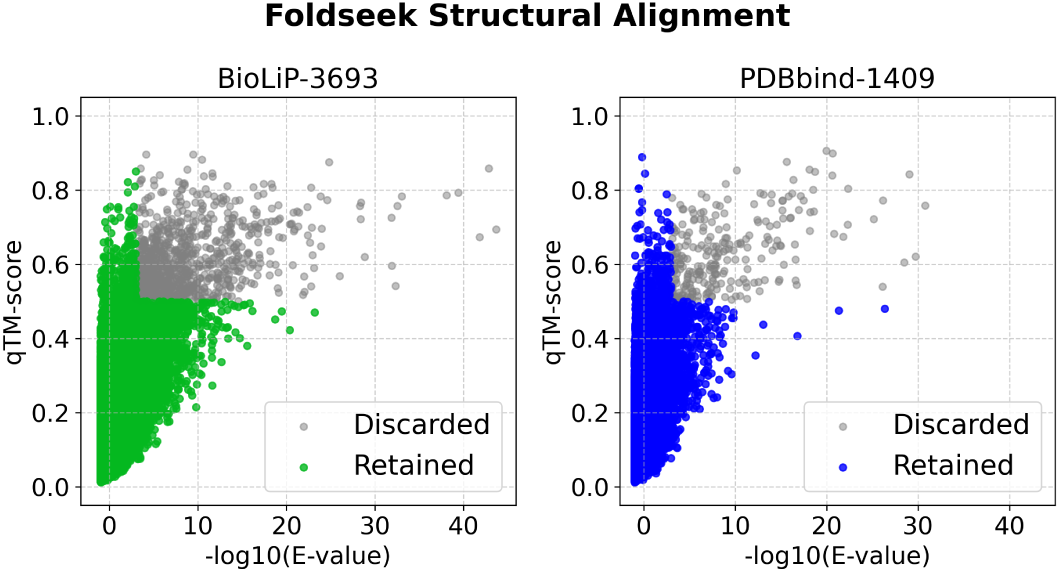
Data Leakage Prevention via Structural Homology Filtering. Foldseek alignments quantifying structural similarity by qTM-score and statistical significance, between pairs from the ZK448 benchmark and the two training sets, BioLiP (left) and PDBbind (right). Colored regions represent retained points after the filtering process.

This procedure yields final, non-redundant training sets of 3693 proteins for BioLiP and 1409 for the PDBbind-derived dataset, with positive instances accounting for 11% and 13% of the total residues, respectively.

We also curate an independent dataset derived from the Protein-Protein Docking Benchmark version 5.0 (**DB5**) [63]. We restrict our selection to DB5 entries featuring exactly two interacting chains, yielding an initial set of 100 dimeric complexes. To guarantee a strictly unbiased evaluation, we apply the dual-stage homology filtering protocol against the training sets. Furthermore, we exclude complexes exhibiting an unbound-to-bound Root Mean Square Deviation (RMSD) greater than 10 Å to prevent severe conformational changes from confounding interface RMSD assessments. This filtering protocol yields a final benchmark set of 12 non-redundant complexes (Table S1).

### 2.2 ESM3 Refinement for Interaction Site Identification

We implement a frozen-weight transfer learning baseline and an end-to-end finetuning approach to adapt the ESM3 model [56] for PPI sites identification, resulting in **ESM3-PPISites**. Relying exclusively on the protein sequence as input, we extract embedding representations from the final hidden layer of the open-weight ESM3 (1.4B), and the large-scale, proprietary version (98B). We classify interface residues by passing representations through a lightweight downstream head, consisting of a single fully connected layer. The network is optimized using a binary cross-entropy loss applied to the resulting classifier logits. To account for the severe class imbalance, the loss incorporates a dynamic positive class weighting scheme. Training was performed using the AdamW optimizer with early stopping based on the validation loss, using a tolerance of 5 epochs without improvement. In the baseline setting, the models were kept frozen and L2 normalization was applied to each residue embedding prior to classification. In this setting, we adopt a learning rate of 5 *·* 10*^−^*^4^ and a batch size of 96. When finetuning the smaller model, gradients are backpropagated through the transformer blocks of the ESM3 model, allowing the representations to adapt to the downstream task. To stabilize training, Layer Normalization was applied across the hidden dimension. For both the PDBbind and BioLiP datasets, we apply a learning rate of 2 *·* 10*^−^*^5^ and a weight decay of 0.05, utilizing a batch size of 1 with gradient accumulation over 4 steps. The larger ESM3 variant was used in baseline mode only, since this model is not publicly available and does not permit end-to-end finetuning. All computations were conducted using NVIDIA DGX A100 and DGX H100 systems.

### 2.3 Translation of Interface Predictions into Docking Restraints

We evaluate the possible application of our model in a protein-protein interaction prediction context by performing HADDOCK experiments [64] on a set of 12 complexes with experimentally determined structures. The performance of our prediction-guided docking is compared against both a blind docking control and a positive control using native-contact restraints through the HADDOCK CAPRIeval protocol, calculating the interface RMSD (iRMSD) relative to the native complex. For the unrestrained **ab initio** runs, we employ a rigid-body sampling of 10,000 models followed by flexible refinement of the best 200 models. While extended sampling ranges were explored, increasing to 15,000 trials provides no performance improvement, and halving to 5,000 trials proved insufficient to sample high quality configurations. This confirms that 10,000 trials represent an optimal balance of computational efficiency and accuracy. The **oracle** upper bound for docking performance is established using a ground-truth reference utilizing native interfacial contacts extracted from the experimental complexes. These interactions are defined as *Cα* pairs on opposing chains within an 8Å distance threshold, and modeled as unambiguous distance restraints with a target of 10.0Å. For this reference run, we generate 100 rigid-body models and refine the top 10 candidates through energy minimization.

To drive the docking process, we implement a **prediction-guided** HADDOCK protocol that maps predicted residue-level interfaces of single proteins onto spatial distance restraints between the interacting partners (Fig. 3). First, we cluster predicted residues into ‘patches’, defined as the groups of predicted residues separated by gaps of at most three positions. We select the two patches per chain with the highest mean predicted probability and systematically evaluate all patch-wise combinations between the two chains. For each pair, the shorter patch (*L*_short_) is centered relative to the longer one (*L*_long_) using an offset *s* = (*L*_long_ *− L*_short_)*/*2. This alignment provides the anchor for our restraints; specifically, to account for registration uncertainties, we map each residue *r*_short_*_,i_* to a three-residue window on the longer patch, spanning the aligned position *r*_long_*_,s_*_+_*_i_* and its flanking neighbors *r*_long_*_,s_*_+_*_i±_*_1_. To accommodate both parallel and anti-parallel binding modes, the shorter patch is evaluated in both forward and reverse orientations. Contacts are then enforced via Ambiguous Interaction Restraints, which use the same distance parameters as the oracle reference. We process each set of constraints through a docking run consisting of 100 rigid-body models, followed by the selection and energy minimization of the best-performing model. We then rely on HADDOCK scoring to rank the obtained binding modes.

**Fig. 3.**
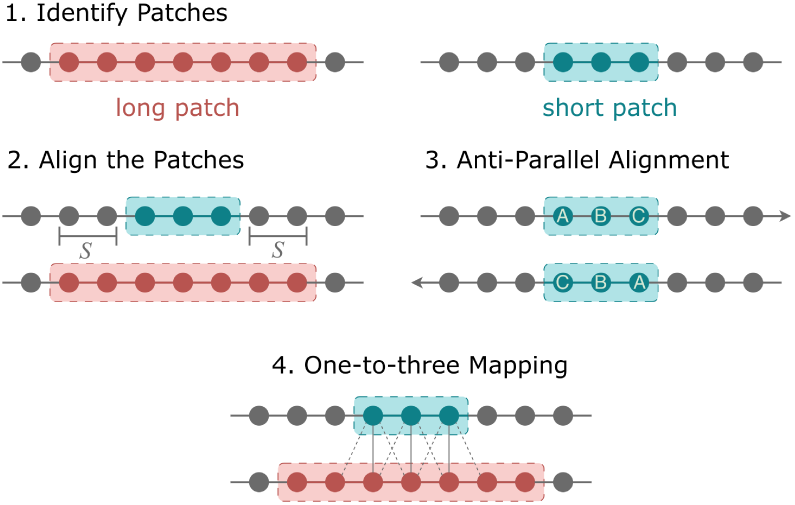
Mapping of predictions onto docking restraints. (1) Patches identified as contiguous predictions (2) aligned by centering the shorter patch (3) both parallel and anti-parallel orientations pairing (4) one-to-three mapping to the longer patch for ambiguous interaction restraints.

## 3 Results

### 3.1 Performance Evaluation

In the context of strongly unbalanced classifications, where positive interface residues are vastly outnumbered by non-interacting residues, we employ a comprehensive suite of metrics. The F1 score isolates success on the critical minority class by disregarding the massive true-negative rate. Complementing this, the MCC provides a symmetric, balanced summary of the entire confusion matrix, ensuring a high score is achieved only if the model performs well on both the minority and majority classes. As illustrated in Fig. 4, all evaluated architectures demonstrate highly competitive predictive capabilities. Even the unoptimized baseline 1.4B parameter model (ESM3-1.4B) achieves F1 scores above 0.60 and MCC values exceeding 0.50, confirming that the pre-trained embeddings already inherently encode substantial structural and functional information. As expected, scaling to the massive 98B parameter model (ESM3-98B) consistently delivers the highest overall performance, pushing the F1 score near 0.70 and the MCC above 0.60. However, supervised finetuning of the smaller model (FT-ESM3-1.4B) significantly narrows this performance gap. This finding is particularly relevant, as it shows that targeted optimization on domain-specific data can robustly compensate for the lower native capacity of smaller models, providing a highly efficient alternative to larger-scale, proprietary architectures. Furthermore, the models exhibit consistent predictive behavior across training datasets derived from distinct sources. Despite a significant disparity in dataset scale (3 693 proteins for BioLiP versus 1 409 for PDBbind), the performance metrics remain highly comparable. While the larger BioLiP dataset yields higher absolute scores, the good performance maintained on the smaller PDBbind set reveals that the model is able to generalize effectively regardless of the specific training data employed.

**Fig. 4.**
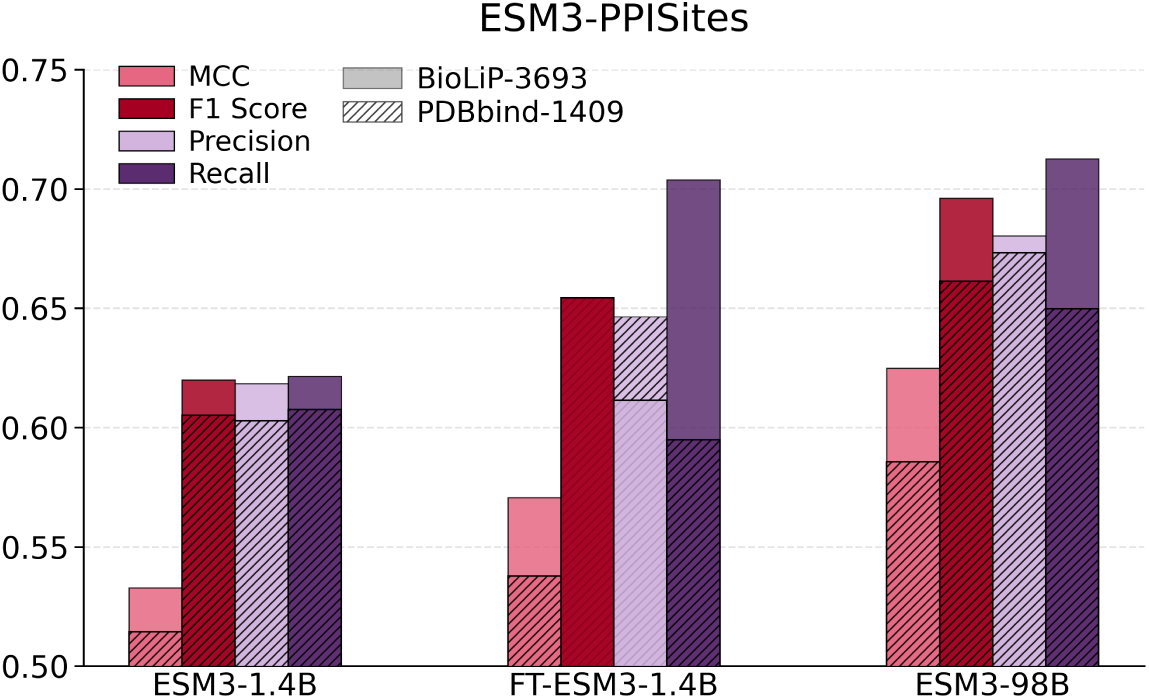
Performance of ESM3-based architectures. Baseline (ESM3-1.4B), scaled (ESM3-98B) and finetuned (FT-ESM3-1.4B) models evaluation on the ZK448 test set (through MCC, F1 Score, Precision, and Recall metrics). Results are stratified by training source, comparing BioLiP-3693 (solid) against PDBbind-1409 (hatched) datasets.

A detailed analysis of the predictive behavior of ESM3-PPISites on the independent ZK448 benchmark, both at the residue and protein levels, focuses on the physicochemical properties of the identified residues and the consistency of the model’s performance across individual protein targets. As illustrated by the log-odds ratio of predicted interactions grouped by amino acid class (Fig. 5A), the model successfully recovers the established signatures of protein-protein interfaces. The predictions demonstrate a marked enrichment for hydrophobic and aromatic residues, which are well known to dominate binding interfaces and serve as energetic hot spots. Conversely, polar and charged residues display lower predicted interaction frequencies, consistent with the biophysical constraints governing protein-protein molecular recognition. Furthermore, the distribution of Relative Solvent Accessibility for the predicted interface residues (Fig. 5B) shows that the model accurately targets surface-exposed regions. The density sharply increases and peaks above the standard 0.2 consensus cutoff for solvent accessibility [65], indicating that the model hardly predicts buried core residues as interaction sites. Remarkably, when quantifying this surface-exposed population, the ratio of predicted interacting residues to the total number of exposed amino acids closely matches the ground-truth proportion of truly interacting residues within the exposed surface (26%). This alignment reveals two critical insights: first, it confirms that the model is not simply a surface-exposure detector that over-predicts accessible sites, but instead remains highly selective; second, it demonstrates that the model preserves the native interface density observed in experimentally resolved complexes. Comparing the F1 score and MCC for each protein in the benchmark set shows that the majority of them achieve scores well above 0.5 in both metrics, with few pathological failures (Fig. 5C). Finally, when comparing the true against the predicted fraction of interacting residues per protein (Fig. 5D), we observe a strong correlation (Pearson r = 0.81), indicating that there is no systematic bias toward over- or under-predicting the interface area.

**Fig. 5.**
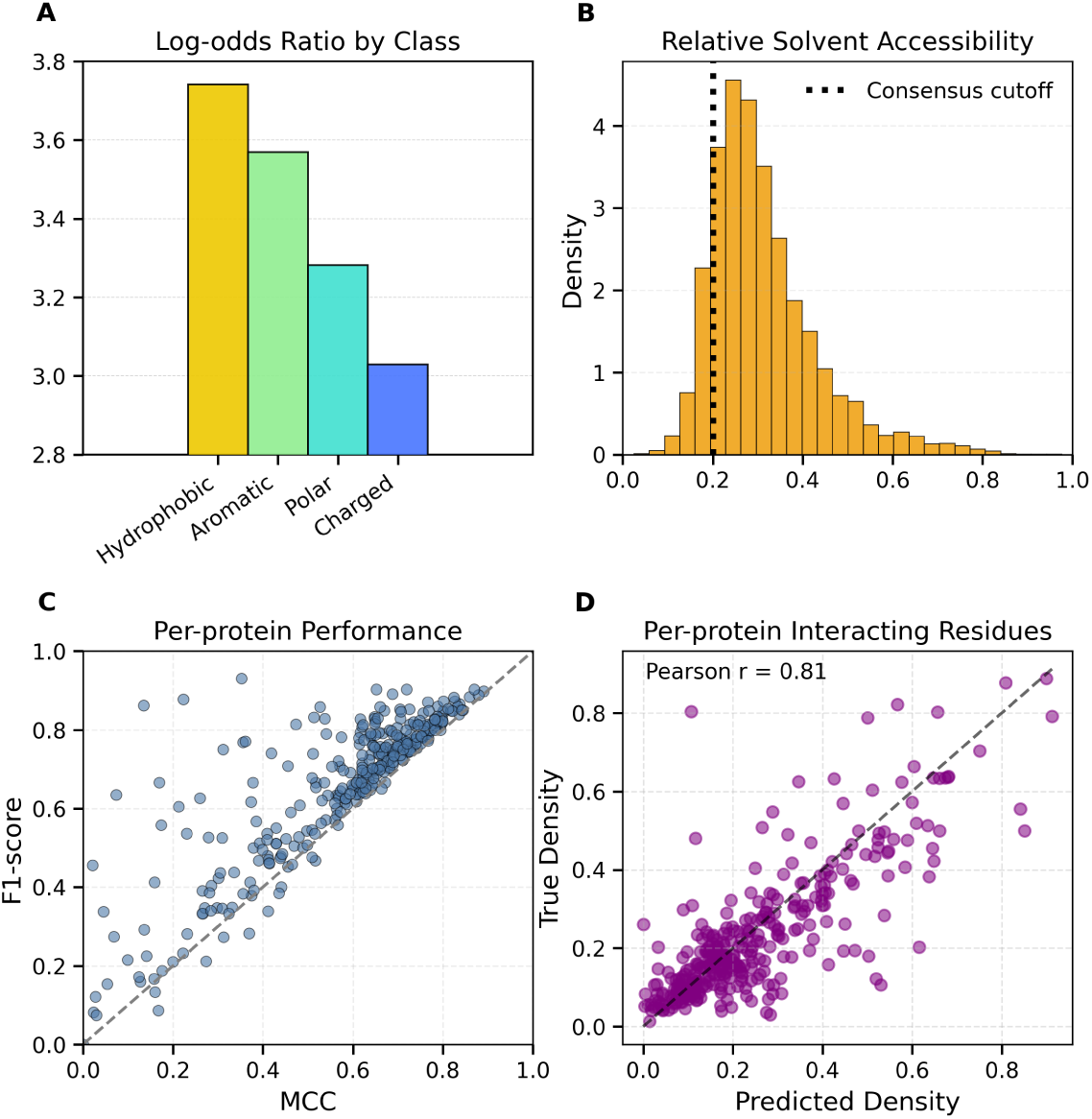
Analysis of predictions. **(A)** Log-odds ratio of predicted interacting residues categorized by biochemical class. **(B)** Density distribution of Relative Solvent Accessibility for predicted interface residues. **(C)** Per-protein performance evaluation. **(D)** Correlation between the true and predicted fraction of interacting residues per protein.

### 3.2 Comparison with Existing Methods

#### 3.2.1 Comprehensive Benchmark Evaluation

We compare ESM3-PPISites with established sequence- and structure-based interface prediction methods. Results for sequence-based methods are taken from a recent study [54], whereas PeSTo and ScanNet are evaluated using the official implementations released by their authors [42, 44]. As reported in Table 1, ESM3-PPISites achieves the best results across all considered metrics. The ESM3-98B-powered variant attains an F1 score of 0.696 and an MCC of 0.625, substantially outperforming the evaluated structure-based methods, ScanNet (F1: 0.390, MCC: 0.215) and PeSTo (F1: 0.317, MCC: 0.144). ESM3-PPISites also outperforms the strongest sequence-based baseline, Seq-InSite, achieving an F1 score of 0.696 versus 0.535 and an MCC of 0.625 versus 0.462. Altogether, these gains demonstrate that ESM3-PPISites provides a substantial improvement over existing predictors, without requiring the structure of the protein at inference time. Overall, its consistent advantage supports the value of multimodal protein representations for residue-level interface prediction.

**Table 1.** Performances on the ZK448 benchmark.

| Model | Ref. | F1 score | MCC | AUC |
| --- | --- | --- | --- | --- |
| *Sequence-based:* |  |  |  |  |
| ESM3-PPISites |  | **0.696** | **0.625** | **0.913** |
| Seq-InSite | [53] | 0.535 | 0.462 | 0.853 |
| PIPENN-EMB | [54] | 0.505 | 0.392 | 0.805 |
| SCRIBER | [19] | 0.333 | 0.230 | 0.715 |
| SSWRF | [22] | 0.287 | 0.178 | 0.687 |
| CRFPPI | [23] | 0.266 | 0.154 | 0.681 |
| LORIS | [18] | 0.263 | 0.151 | 0.656 |
| SPRINGS | [17] | 0.229 | 0.111 | 0.625 |
| PSIVER | [26] | 0.191 | 0.066 | 0.581 |
| SPRINT | [21] | 0.183 | 0.057 | 0.570 |
| SPPIDER | [20] | 0.198 | 0.071 | 0.517 |
| EnsemPPIS | [52] | 0.385 | 0.291 | 0.770 |
| DELPHI | [28] | 0.364 | 0.278 | 0.746 |
| PIPENN | [29] | 0.339 | 0.249 | 0.755 |
| *Structure-based:* |  |  |  |  |
| PeSTo | [44] | 0.317 | 0.144 | 0.670 |
| ScanNet | [42] | 0.390 | 0.215 | 0.673 |

#### 3.2.2 Evaluation on Restricted Training Regime

To test generalization under severe data scarcity, we train and test ESM3-PPISites using datasets from RGLLA-PPIS [40], a recently published framework that relies on an intricate pipeline chaining AlphaFold3 structures, equivariant graph neural networks, and retrieval-augmented language priors. Operating under their exact restricted training protocol and evaluating on the independent test315 and test71 partitions, these data-limited configurations expose a steep generalization bottleneck for all models (Table 2). On the test315 benchmark, the multi-component architecture of RGLLA-PPIS maintains an advantage, achieving an F1 score of 0.49 and an MCC of 0.39, compared to an F1 score of 0.45 and an MCC of 0.35 for ESM3-PPISites. However, on the independent test71 split ESM3-PPISites outperforms all competing baselines, surpassing RGLLA-PPIS across all metrics.

**Table 2.**
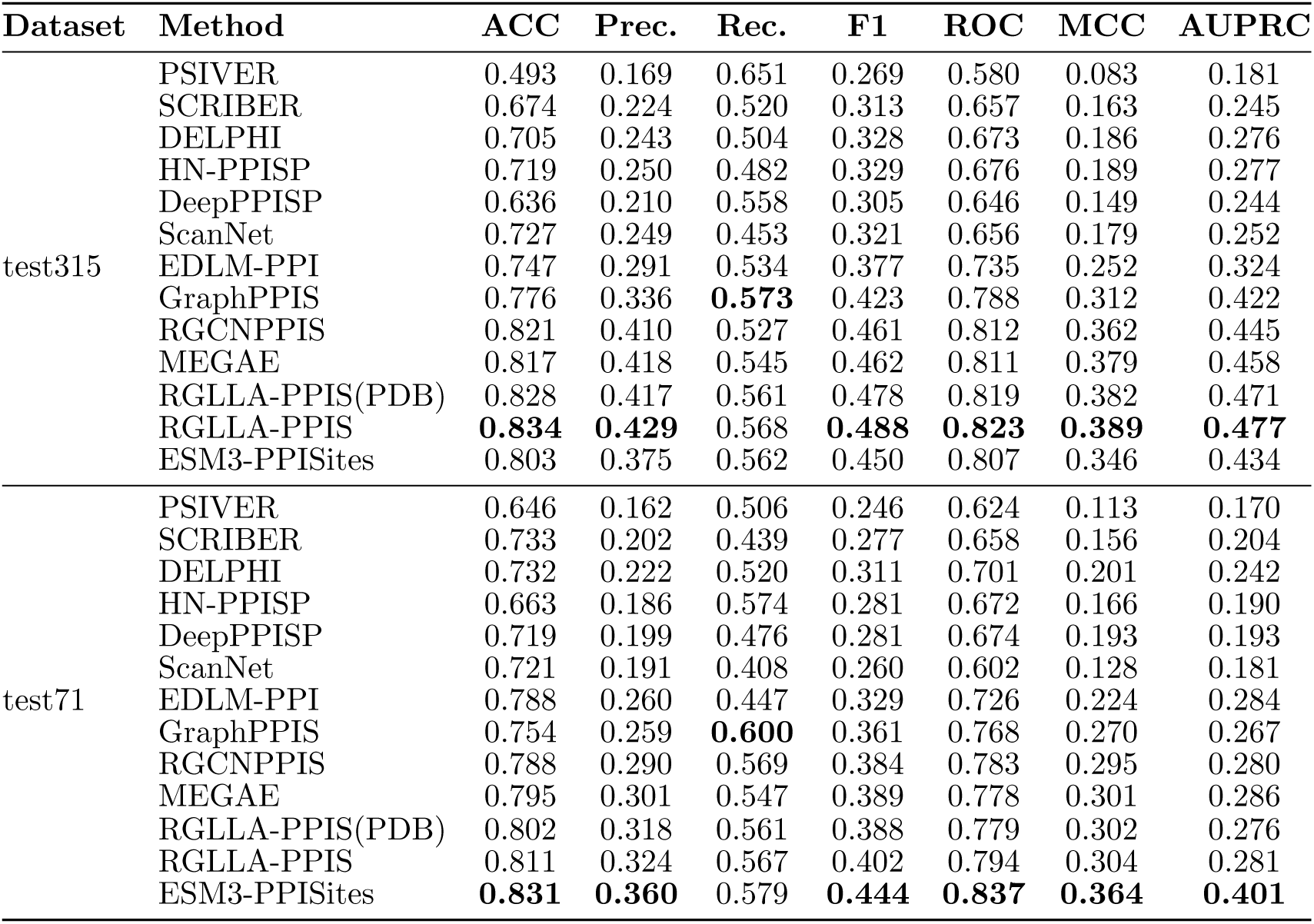
Comparative performance on test315 and test71 under the train324 regime of RGLLA-PPIS.

#### 3.2.3 Comparison with AlphaFold3 on Challenging Complexes

A direct comparison between ESM3-PPISites and AlphaFold3 (AF3) requires careful consideration, as the two approaches address different tasks. ESM3-PPISites is finetuned specifically for residue-level protein–protein interface site classification, whereas AF3 is a biomolecular structure prediction model. Moreover, establishing equivalent control over training data overlap is particularly challenging for AF3, whose training data and potential exposure to homologous proteins cannot be controlled in the same way as for a task-specific model trained using explicitly defined data splits. This distinction is important when interpreting performance differences, since structural similarity to training complexes can influence AF3’s ability to predict protein–protein interactions. Acknowledging these limitations, we select two sets of protein complexes for comparison. The first set comprises protein complexes from the 16th Critical Assessment of Structure Prediction (CASP16) [66], a community-wide blind assessment in which predictions are submitted before the corresponding experimental structures are disclosed. This setting provides a particularly challenging benchmark, as the targets are not yet structurally characterized at the time of prediction and many oligomeric complexes remain difficult to model accurately, even for state-of-the-art methods. Despite this, it is not guaranteed that AF3 has not encountered proteins related to these targets during training; potential leakage therefore remains an issue. The second set comprised the 13 antibody–antigen complexes with the lowest DockQ scores in a recent systematic evaluation of AF3, representing particularly challenging cases for complex modeling. We restrict the analysis to complexes with fewer than 1,100 residues across their constituent chains and with complete experimentally resolved interfaces. Across the CASP16 complexes and the selected low-DockQ antibody–antigen complexes, AlphaFold3 generally achieved higher overall F1 scores (Fig.6) and precision across most targets, whereas ESM3-PPISites often showed competitive or higher recall (Figures S1 and S2). These effects should be interpreted in light of the distinct objectives of the two approaches: AF3 predicts the structure of a specific biomolecular complex at the atomic level, from which interface residues are identified based on their spatial arrangement in that complex, whereas ESM3-PPISites classifies residues based on interface patterns learned through supervised training across interaction contexts, irrespective of the specific partner under evaluation. Notably, on the most challenging low-DockQ antibody–antigen complexes, failing AF3 structural predictions were often accompanied by low ESM3-PPISites performance. This observation suggests ESM3-PPISites does not overcome the limitations associated with inherently challenging systems. Overall, AlphaFold3 demonstrates a general performance advantage across the evaluated datasets, yet ESM3-PPISites remains competitive, frequently approaching AF3’s level of accuracy. For the CASP16 targets, AF3 exhibits a marked advantage on multi-chain complexes such as 9C3C, spanning components of the dystrophin-associated protein complex (beta-dystroglycan, dystrophin, dystrobrevin, alpha-sarcoglycan, and sarcospan), as well as chains D and E of 9NB9, corresponding to skeletal muscle alpha-actin and deoxyribonuclease-1, respectively. ESM3-PPISites achieves a higher F1 score for chain B of 9CBN (Fig. 7), which corresponds to the kappa chain of the HAstV1-neutralizing Fab 2B4. In the antibody–antigen benchmark, AF3 outperforms ESM3-PPISites on several targets, though the magnitude of this advantage varies by chain. This performance gap is particularly remarkable for 8DWX, where AF3 maintains superior performance across multiple chains belonging to the E1 and E2 glycoproteins and the capsid protein. Nevertheless, ESM3-PPISites demonstrates clear advantages on selected nanobody and antibody chains, such as chain B of 8U3S, chain E of 8BF4, and chain B of 8SBB. [67].

**Fig. 6.**
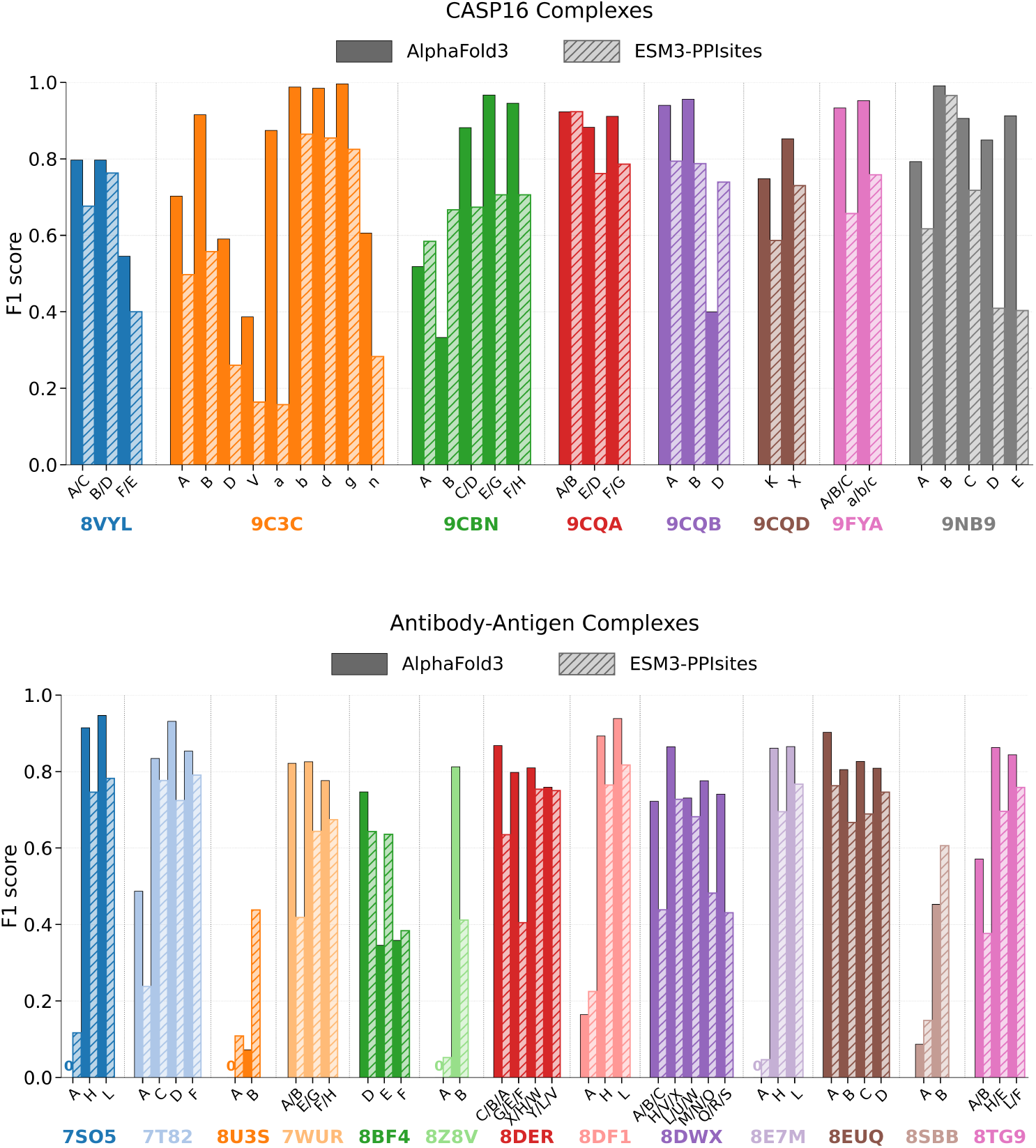
Comparison of interface residue classification performance between AlphaFold3 (solid) and ESM3-PPISites (hatched). F1 score for individual chains across CASP16 targets (Top) and challenging anti-body–antigen complexes (Bottom).

**Fig. 7.**
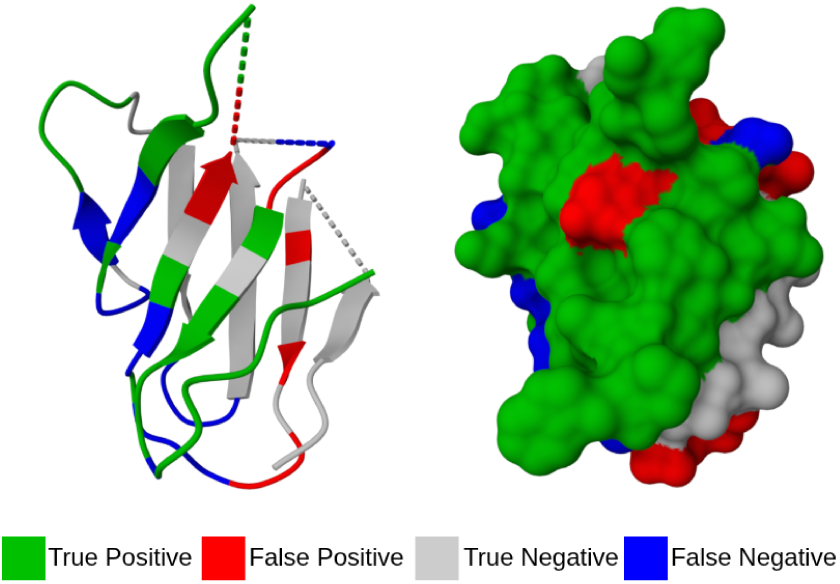
Residue-level classification of ESM3-PPISites on the *κ* chain of the HAstV1-neutralizing Fab 2B4 (PDB: 9CBN, chain B), represented in cartoon (left) and surface (right) modes.

### 3.3 Guiding Docking with ESM3-PPISites Restraints

We integrate the ESM3-PPISites output as spatial restraints in HADDOCK to reduce the conformational search space and guide complex assembly. We measure the interface Root Mean Square Deviation (iRMSD) and DockQ score relative to the experimental structures of the complexes and compare the results of the prediction-guided protocol against a computationally intensive *ab initio* blind docking baseline and an *oracle* reference utilizing true native contacts. The *oracle* set-up serves as a positive control, and the procedure consistently yields configurations with an iRMSD below 5Å and high DockQ scores across evaluated complexes. As illustrated in Fig. 8, showing the relationship between iRMSD and DockQ score, we consider the 10 lowest HADDOCK score configurations generated by both the ESM3-PPISites-guided protocol and the *ab initio* baseline. We define successful docking runs as those sampling at least one configuration with an iRMSD below a 10Å cutoff. Under these criteria, the ESM3-PPISites protocol achieves a high success rate, yielding low iRMSD models for 9 out of the 12 evaluated complexes, substantially outperforming the ab initio baseline, which successfully samples acceptable configurations for only four complexes (half of which yield zero DockQ scores). Remarkably, our prediction-guided approach achieves a high-resolution iRMSD lower than 5Å in 5 successful runs (1Z5Y, 1ZHH, 2YVJ, 2AJF, 3PC8), and these highly accurate configurations rank within the top three scored models. In Figure S3 we extend this docking evaluation to bound experimental structures and AlphaFold3-derived structures, which naturally act as an oracle-level baseline achieving higher overall performance.

**Fig. 8.**
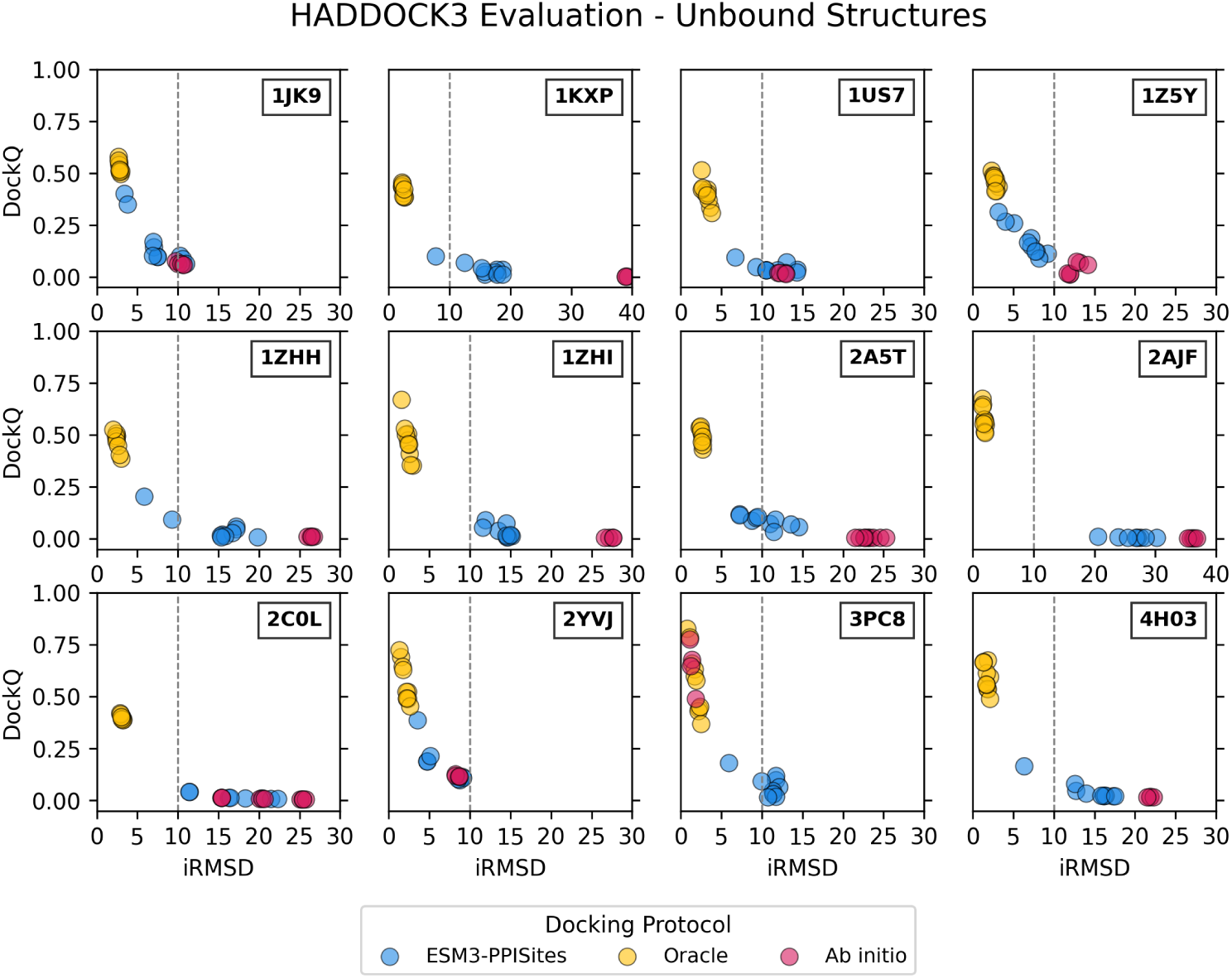
Comparison of docking performance. iRMSD vs DockQ of top ten ranked poses from ESM3-PPISites-guided, oracle and ab initio protocols. The dashed line indicates the 10Å iRMSD successful docking cutoff.

Beyond improving the quality of the resulting structural ensembles, the integration of ESM3-PPISites predictions significantly accelerates the docking process. Fig. 9 details the computational runtime across the three benchmarking strategies carried out on a 64-core AMD EPYC 9374F node (32 cores per job). The unguided ab initio protocol exhibits massive computational overhead, requiring between 20 and 90 minutes per complex due to the broad rigid-body sampling required to blindly explore the interaction landscape. In contrast, restricting the search space using our predicted patches drastically truncates the required runtime to less than 20 minutes for almost all evaluated complexes, mirroring the optimal computational efficiency of the oracle reference.

**Fig. 9.**
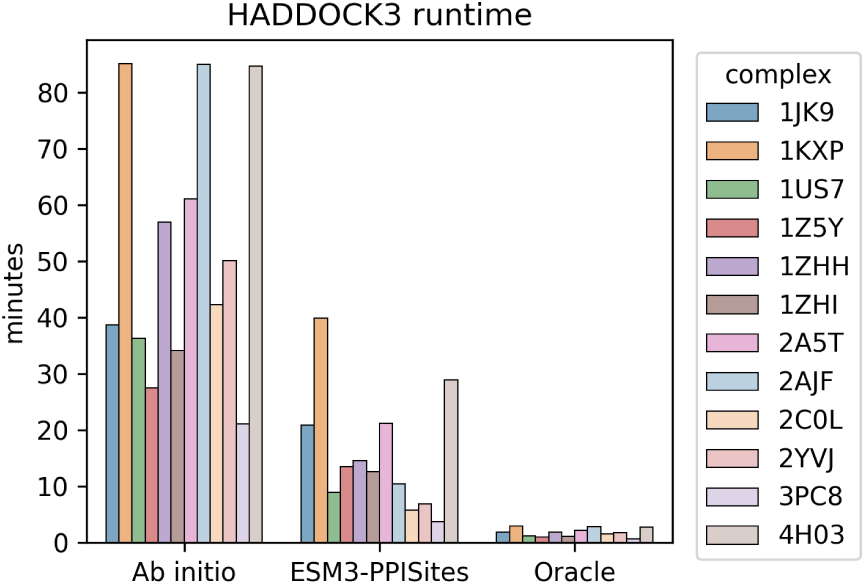
HADDOCK runtimes across protocols and complexes.

Taken together, these findings highlight that translating our ESM3-PPISites predictions into spatial restraints for Haddock simultaneously enhances the resolution of the predicted macromolecular complexes and eliminates the severe computational bottlenecks associated with traditional blind docking.

## 4 Discussion

Accurate identification of protein–protein interaction interfaces is essential for understanding molecular recognition and guiding the characterization and design of protein complexes. Here, we introduce ESM3-PPISites, a residue-level interface classification model developed through supervised finetuning of the multimodal ESM3 PLM. The model predicts interface residues directly from individual protein sequences, without requiring the structure of the interacting complex at inference time. Model evaluation incorporates stringent sequence- and structure-based redundancy filtering to assess generalization beyond closely related proteins. finetuning the open-weight ESM3-1.4B model substantially narrows the performance gap with the proprietary ESM3-98B variant, highlighting the potential of task-specific adaptation to make multimodal protein representations more accessible for residue-level interface prediction. ESM3-PPISites captures key physicochemical properties of protein–protein interfaces, preferentially identifying hydrophobic and aromatic residues while rarely assigning interface labels to buried residues. Its predictions are not driven by surface exposure alone: the proportion of predicted interface residues among exposed residues closely matches the experimentally observed fraction. Moreover, the strong correlation between predicted and observed interface fractions indicates that the model reflects differences in interface extent across proteins. Under a restricted training regime, ESM3-PPISites maintains strong performance, ranking among the best-performing models in one evaluation and outperforming all competing baselines across metrics in another. Moreover, while AlphaFold3 generally yields superior F1 scores and precision across the CASP16 and antibody–antigen benchmarks, ESM3-PPISites remains highly competitive on the majority of targets. Notably, both approaches exhibit concurrent performance degradations on specific target subsets, implying that these shared failure modes stem from inherent structural or physicochemical challenges common to modern interface prediction paradigms rather than model-specific biases. We showcase the practical utility of the model to accelerate PPI modeling by incorporating predicted interface residues as interaction restraints in HADDOCK. For the evaluated complexes, these restraints markedly improve docking success relative to blind docking while drastically reducing computational time.

A primary limitation of the method is that it identifies residues likely to participate in protein interfaces without explicitly determining interaction partner compatibility, binding geometry, or binding specificity. Future work will therefore extend it by integrating interface predictions with partner identification and structural modeling, aiming to connect residue-level propensities with fine-grained structural characterization. In conclusion, ESM3-PPISites demonstrates that multimodal protein language model representations can be effectively adapted for residue-level PPI prediction. By pairing strong predictive accuracy with computational efficiency, this approach establishes a reliable and accessible tool for large-scale interface annotation.

## 5 Availability of Data and Software

The code used in this work is publicly available in the GitHub repository: https://github.com/RitAreaSciencePark/ESM3-PPISites. ESM3 representations and PDB files are stored in Zenodo at https://doi.org/10.5281/zenodo.20366947. The trained model is accessible via Hugging Face at https://doi.org/10.57967/hf/10993. To facilitate ease of use, a web-based inference dashboard is also provided, enabling real-time predictions without local environment configuration: https://huggingface.co/spaces/area-science-park/esm3-ppisites.

## 6 Authors’ Contribution

Y.G. developed and optimized the finetuning pipeline for the ESM3-1.4B model, implemented the Foldseek-based data curation strategy, and managed the GitHub repository. E.N.V.G. developed the finetuning pipeline of the ESM3-1.4B and ESM2 models (not shown), the extraction of representations from the ESM3-98B model, and the Hugging Face deployment. S.S. carried out prediction analyses on the ZK448 test set, performed the PeSTo model evaluation and supported data curation. D.D. curated and optimized the code for baseline ESM3, contributed to predictions analysis, performed the ScanNet model evaluation and managed the Zenodo repository. A.F. assembled the PDBbind-derived dataset, performed data curation, and contributed to prediction analysis on the ZK448 test set. F.C. supervised and coordinated the study and led the writing of the manuscript. All authors contributed to the manuscript writing and revision.

## 7 Acknowledgments and Funding

The authors acknowledge the Area Science Park supercomputing platform ORFEO made available for conducting the research reported in this article and the technical support from the team of the Laboratory of Data Engineering. Y.G. is supported by the Programma Nazionale della Ricerca (PNR) grant J95F21002830001 with the title ‘FAIR-by-design’. E.N.V.G. was supported by the project PON ‘BIO Open Lab (BOL) - Rafforzamento del capitale umano’ (CUP J72F20000940007). S.S. is supported by I-HUB FVG - INNOVATION HUB FVG (CUP J23B24000030002). D.D. is supported by Progetto ‘QuB - Quantum Behavior in Biological Functions’ (CUP J95F21002820001) financed by ‘Fondo ordinario per gli enti e le istituzioni di ricerca’ (FOE) 2021.

## 9 Use of Large Language Models

Large language models were used in an open-access setting to provide support for language editing. Their use was restricted to auxiliary tasks such as improving text clarity, without any involvement in the design and execution of this work.

## Supporting Information

**Table S1:**
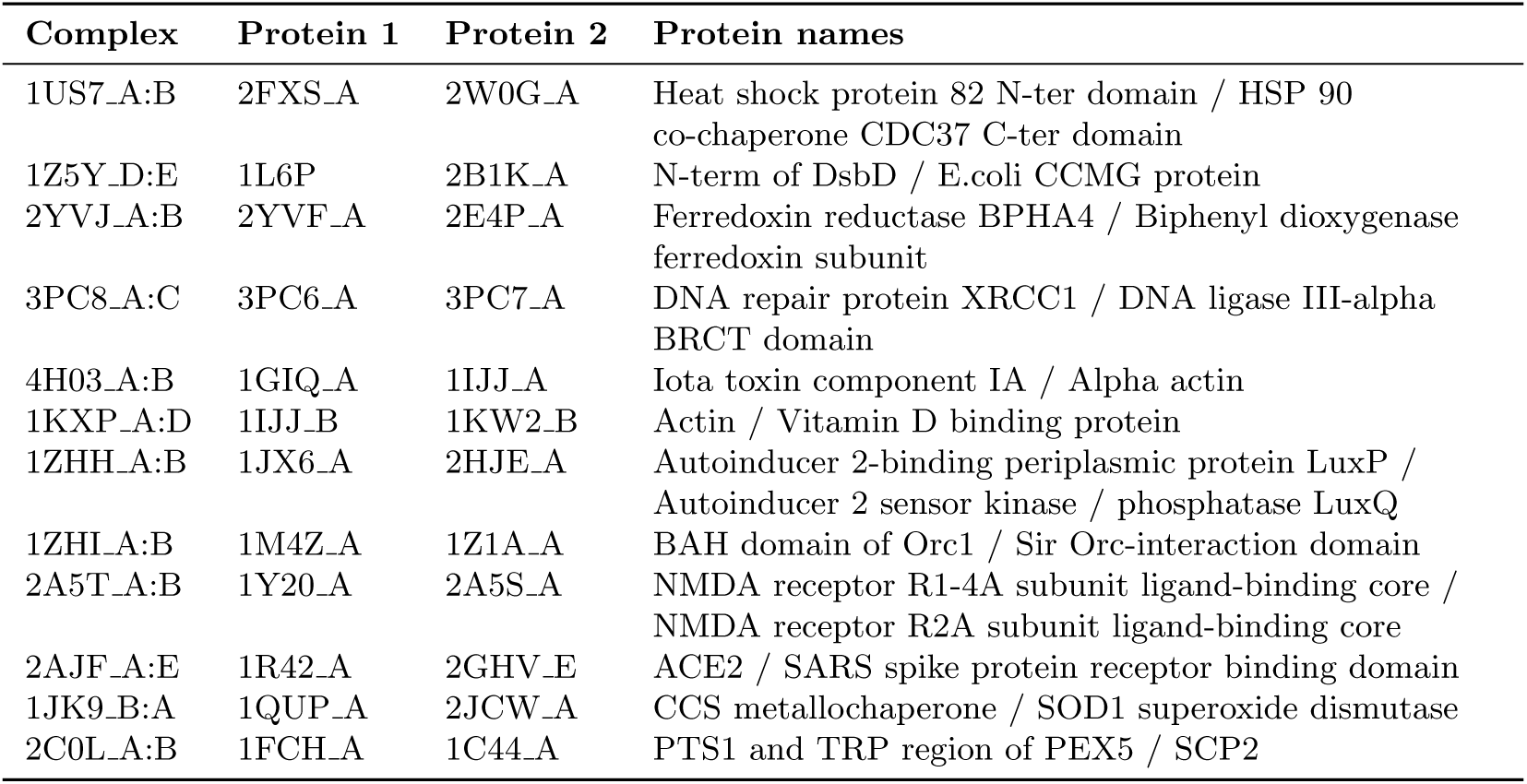
Protein complexes for leakage-free evaluation obtained from the DB5 database.

**Figure S1:**
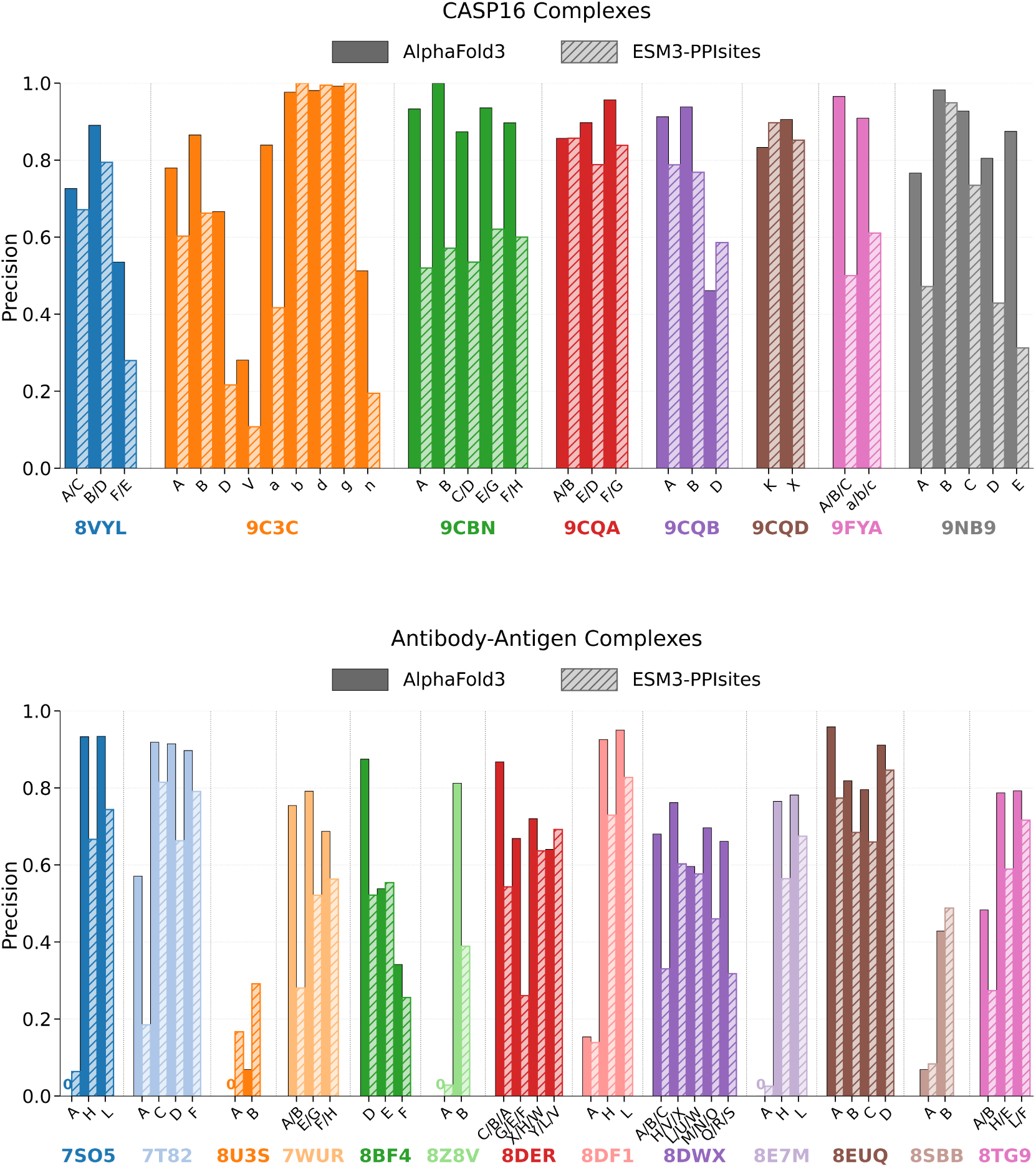
Comparison of interface residue classification performance between AlphaFold3 (solid) and ESM3-PPISites (hatched). Precision for individual chains across CASP16 targets (Top) and challenging antibody–antigen complexes (Bottom).

**Figure S2:**
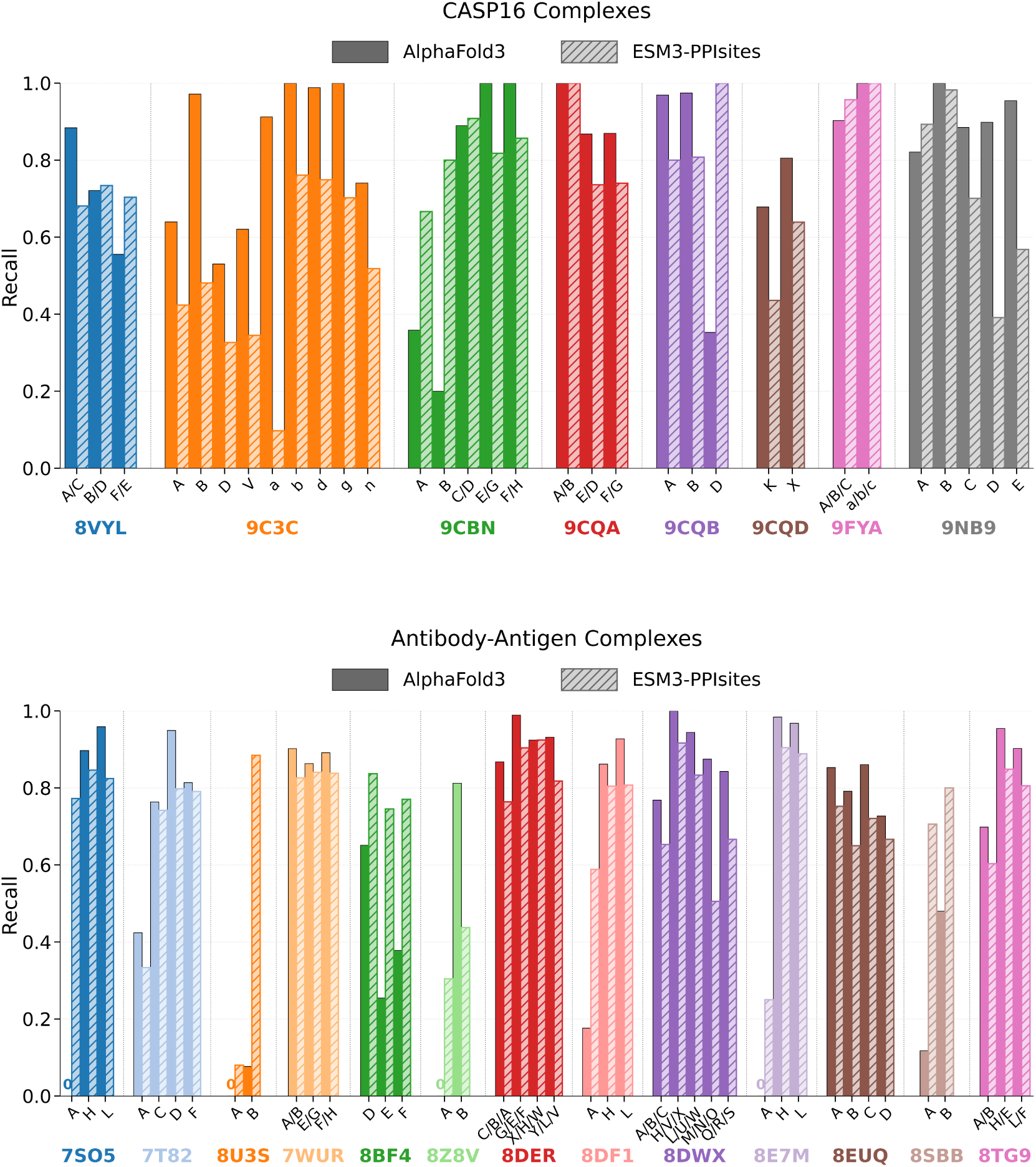
Comparison of interface residue classification performance between AlphaFold3 (solid) and ESM3-PPISites (hatched). Recall for individual chains across CASP16 targets (Top) and challenging antibody–antigen complexes (Bottom).

**Figure S3:**
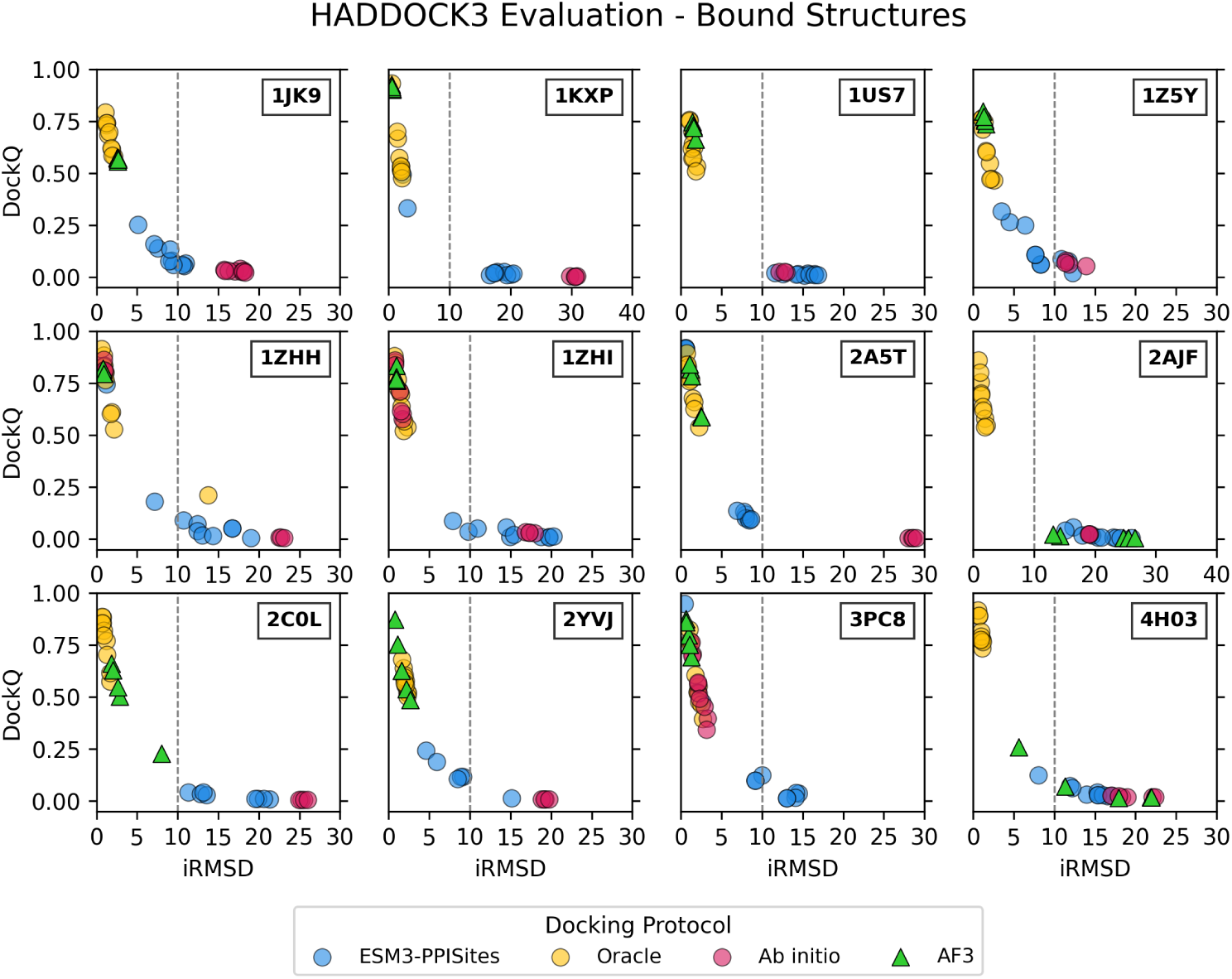
Comparison of docking performance. iRMSD vs DockQ of top ten ranked poses from ESM3-PPISites-guided, oracle and ab initio protocols, together with 5 AlphaFold3 models. The dashed line indicates the iRMSD 10Å cutoff used to define successful docking.

